# Stress resilience is established during development and is regulated by complement factors

**DOI:** 10.1101/2022.01.31.478444

**Authors:** Amrutha Swaminathan, Michael Gliksberg, Savani Anbalagan, Noa Wigoda, Gil Levkowitz

## Abstract

Individuals in a population respond differently to stressful situations. While resilient individuals recover efficiently, others are susceptible to the same stressors. However, it remains challenging to identify resilience in mammalian embryos to determine if stress resilience is established as a trait during development or acquired later in life. Using a new behavioural paradigm in zebrafish larvae, we show that resilience is a trait that is determined and exhibited early in life. Resilient and susceptible individuals retained these traits throughout life and passed them on to the next generation. Resilient larvae showed higher expression of resilience-associated genes and larvae lacking neuropeptide Y and miR218 were significantly under-represented in the resilient population. Unbiased transcriptome analysis revealed that multiple factors of the innate immune complement cascade were downregulated in resilient larvae in response to stressors. Pharmacological inhibition and genetic knockouts of critical complement factors led to an increase in resilience. We conclude that resilience is established early during development as a stable trait, and that neuropeptides and the complement pathway play positive and negative roles in determining resilience respectively.

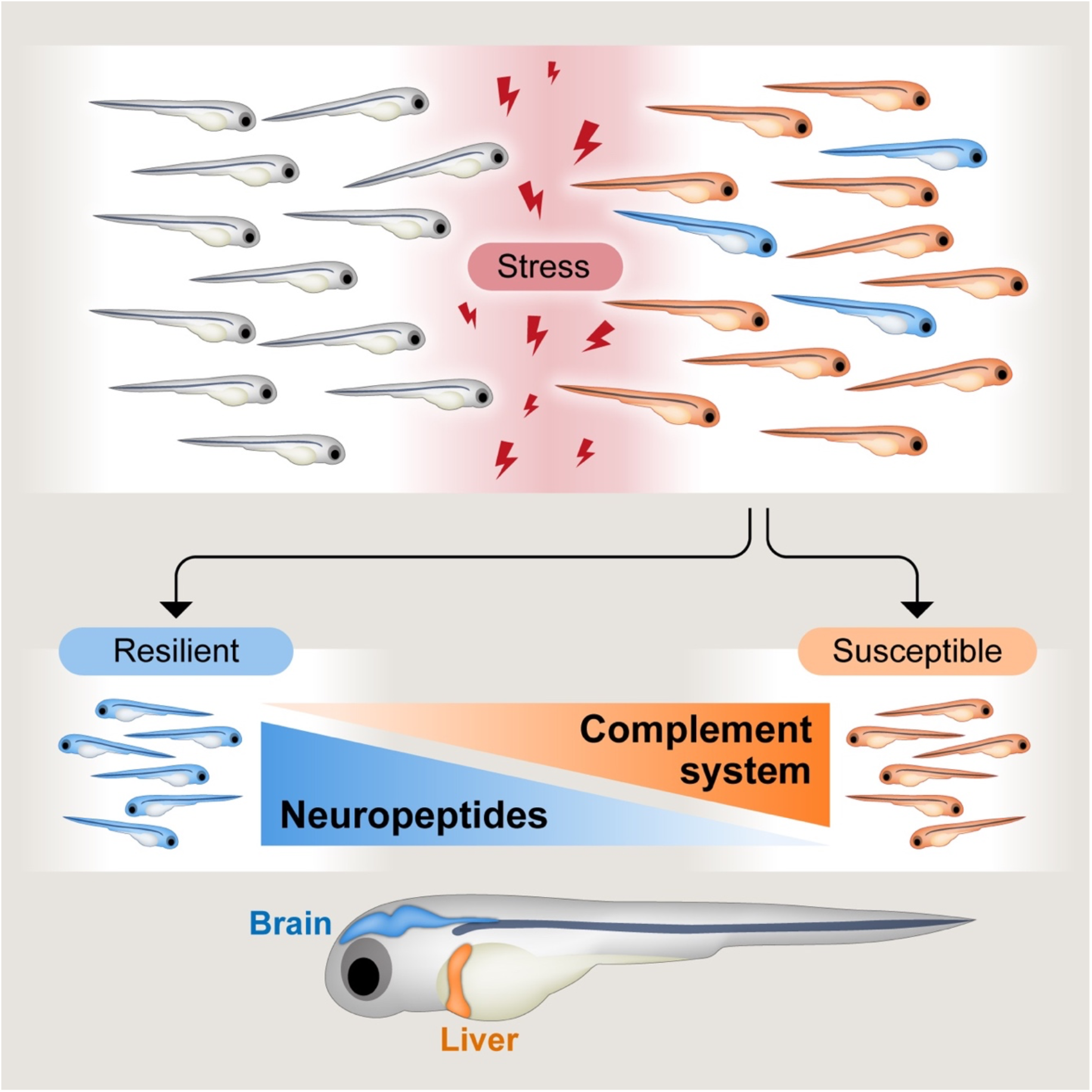

Why some individual individuals recover better than others from stressful situations is unclear. We show that resilience to stress is established during zebrafish development as a stable and heritable trait. Resilience is augmented by brain-derived neuropeptides and attenuated by innate immune complement factors specifically expressed in the liver.

**Highlights:**

- Wildtype zebrafish larvae show differences in their dynamics of recovering from stress, with some being more resilient than others.
- Resilience is a stable and heritable trait.
- Resilient fish show specific stress-responsive transcriptional changes.
- Neuropeptide Y and miRNA218 positively affect resilience, while innate immune complement factors attenuate resilience.

## Introduction

Stress is a systemic response to what an organism perceives as threat, and is associated with changes in physiology and behavior (*1*). Individuals within a population respond differently to stressful events, with resilient individuals being better able to rebound quicker and efficiently from stressful situations. Susceptible individuals also have a higher likelihood to develop stress-related disorders like anxiety, depression and post-traumatic stress disorder (PTSD) (*2, 3*). Stress resilience has been identified and studied in various organisms, and studies using humans or rodents show that both genetic and environmental factors influence resilience/susceptibility tendencies (*3*). Extrinsic factors that influence resilience/susceptibility are: early life adversity and nutrition, gut microbiota, age and gender (*3*). Recently, some intrinsic factors, such as specific brain areas, neural circuits, genetic factors and signaling pathways have been associated with resilience (*2–4*). Examples of prominent resilience-associated factors are neuropeptide Y (NPY), corticotropin-releasing hormone (CRH), brain-derived neurotropic factor (BDNF), FKBP5 and miRNA218 (*4, 5*).

Development plays a major role in determining behaviour later in life and it is postulated that neurodevelopmental impairments contribute to maladaptive stress response in the adult (*6, 7*). However, the assessment of stress resilience in mammals cannot be performed in embryos or in newborn animals. The common paradigm of stress resilience in rodents is assessment of rebound from chronic social defeat (*8*), and studies in human are mainly retrospective, using postmortem genetic studies from heterogenous populations (*9*). Using these approaches, it is not possible to determine if stress resilience is established as a trait during development or whether it is acquired following exposure to environmental factors and/or individuals’ life experience. Moreover, parental nurture during pregnancy and early life have a major impact on behaviour and present additional confounding factors in determining the specification of resilient or susceptible traits.

In this study, we aimed at understanding the developmental underpinnings of individual variability in stress resilience using zebrafish whose fundamental stress-responsive physiological mechanisms are similar to mammals. We developed a robust assay to distinguish between resilient and susceptible individuals during development, based on their dynamics of recovering from stress. We demonstrate that resilience is established during embryonic development as a stable life-long and heritable trait. Finally, we found that resilience is modulated by resilience-associated neuropeptides and members of the component pathway, a crucial component of the innate immune system.

## Results

### Variability in recovery from stress is observed early during development

Zebrafish are vertebrates whose embryos undergo external fertilization and development, and their highly conserved and their robust stress-induced behavioural and hormonal responses begin to function early during development (*10*). Since stress resilience is the ability to rebound effectively from stressful situations, we have devised a high-throughput platform in which we subject zebrafish larvae to major stressors, at 6 days post-fertilization (dpf), and thereafter quantify individual ability to rebound from the stressful event. Two routinely used major stressors in zebrafish larvae are physical handling (i.e. netting) and osmotic challenge, both were shown to induce a classical stress-induced cortisol response (*11*). Following either of these major stressors, we recorded larval locomotion in a 96-well plate under a mildly stressful condition, i.e. novel environment in the dark (Figure 1A).

**Figure 1:**
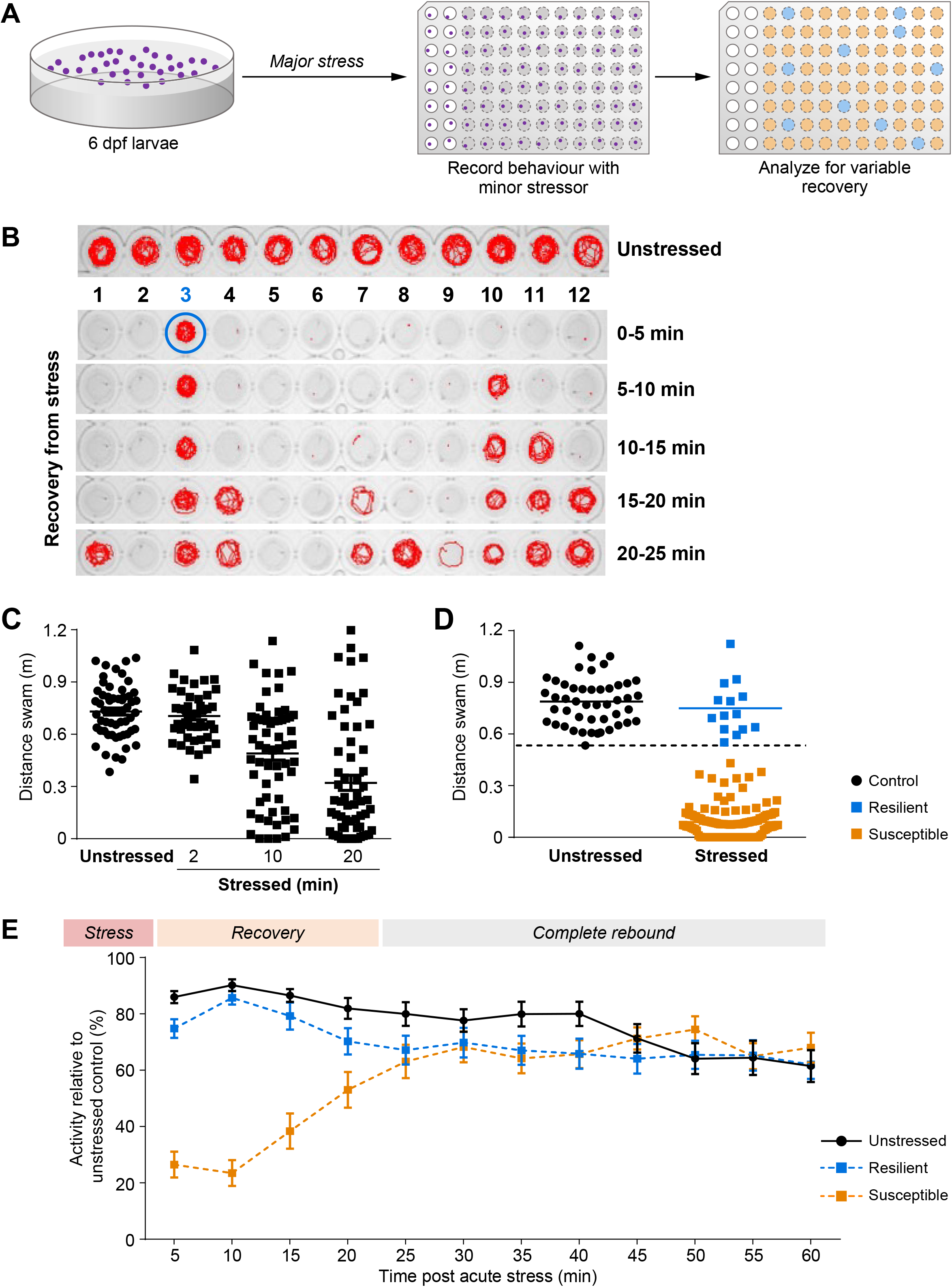
Individual variability in stress rebound in zebrafish larvae. (**A**) Scheme detailing the two-step resilience paradigm. In the first stage, 6 dpf larvae were subjected to a major stressor or left unstressed and each larva was transferred into single wells of a 96-well plate. In the second stage, the recovery of stressed (grey wells) and unstressed (white wells) larvae was monitored under a mild stressful challenge (dark novel environment) and subsequently analyzed. (**B**) Locomotive traces representing the spectrum of behaviours in the resilience paradigm. Unstressed larvae show explorative behaviour, while larvae that underwent major stressful challenge prior to recording show different dynamics of recovery. Lanes 1-12 represent single larva each, and snapshots of their recovery over time is depicted. While some larvae recover very efficiently (lane 3-resilient), others take longer to recover. (**C, D**) Dot plots representing the distance swam during the first 5 minutes of the recovery phase of larvae that were left unstressed or pre-subjected to a major stressor for increasing periods of time. Fast recoverers (resilient-blue) and slow recoverers (susceptible-yellow) individuals in panel D are separated by a dashed line, which represents a statistically calculated threshold. This threshold was determined by calculating the data point where the combined value of Fisher’s exact p value and Cohen’s D value comparing the two archetypes is most significant indicating maximal difference (for details see ‘*Materials and Methods’*). The proportion of the resilient population decreased with increasing time of stress (C). The two groups were best separated after 20 minutes of stress (D). (**E**) Temporal analysis of recovery from a prior stressful challenge of 20 minutes. The resilient group shows behaviour like unstressed control right from the beginning of the task, while the susceptible group recovers more gradually. The difference between the behaviour of these groups is most evident in the first few minutes following stress (variable recovery phase), following which the behaviour of the three groups is comparable (complete rebound phase). Data presented as mean±SEM, **** p<0.0001.

We found that the ability of wildtype individuals to rebound from a stressful event varies within the population in terms of differences in their dynamics of recovery from the stress (Figure 1B). We observed increased individual variability in stress recovery following increasing durations of the stressful challenge (Figure 1C). Thus, with higher intensity of stress, a smaller proportion of the larvae was able to rebound from the stressor (Figure 1C). Hence, to ensure robust differentiation of the resilient and susceptible groups, we chose to perform 20 minutes osmotic stress for further experiments. As a population, the larvae were observed to split in two major archetypes: fast recoverers, termed resilient, whose rebound from the stressful-challenge similar to unstressed controls, and slow recoverers, termed susceptible, which take significantly longer time to rebound from the same major stressor (Figure 1D, 1E). Resilient/susceptible archetypes were classified based on their locomotion in the recovery phase using a statistical thresholding method combining two tools: Cohen’s d effect size and Fisher’s exact test analyzing contingency tables to determine the threshold that best separates the two archetypes (see ‘*Methods’* section).

Since the differences were most pronounced in the beginning of the test, and afterwards all larvae recovered, we chose to analyze the behaviour of the first five minutes that best differentiate between the two archetypes (Figure 1D, 1E). Under these conditions, resilient and susceptible populations were significantly different, with the resilient behaving similar to the unstressed control group (Figure 1D; *p<0.0001*, unstressed n=47, stressed n=144. Kruskal-Wallis test; *p<0.0001* unstressed control and resilient vs. susceptible, *not significant* unstressed control vs. resilient by Dunn’s multiple comparisons test). Using this stress paradigm, we consistently observed that about 10-20% of wildtype populations were defined as resilient (Figure 1D). Susceptible larvae showed significantly differences in stress recovery dynamics compared to the unstressed control and resilient fish (Figure 1E; *p<0.0001*, n=30/group by 2-way ANOVA; *p<0.0001* control vs. susceptible, *p<0.01* resilient vs. susceptible, by Tukey’s post-hoc multiple comparisons test). The recording was performed for an hour to ensure that the susceptible larvae eventually recovered with time, indicating that the observed behavioural differences are related to the unique stress responses of the archetypes (Figure 1E).

In order to analyze whether our larval resilience paradigm was independent of the wildtype strain (AB) used, we performed the assay on another wildtype strain (TL) and found that the archetypes were observed reproducibly in the TL strain (Figure S1A; p*<0.0001*, n=48 unstressed/48 stressed by Kruskal-Wallis test; *p<0.0001* unstressed control and resilient vs. susceptible by Dunn’s multiple comparisons test). Further, when we used a different stressor, i.e., physical handling (netting), we observed similar variability in the stress rebound in the wildtype population (Figure S1B; *p<0.0001*, n=24 unstressed/72 stressed by Kruskal-Wallis test; *p<0.0001* unstressed control and resilient vs. susceptible by Dunn’s multiple comparisons test). We also confirmed that the differences between the resilience and susceptible fish were not due to differences in basal locomotion (Figure S1C *p<0.0001*, n=24 by Kruskal-Wallis test; *p<0.0001* susceptible before vs. after by Dunn’s multiple comparisons test) or due to variable developmental rates in the population (Figure S1D; *p=0.1290*, n=20/20 by Mann-Whitney test).

These results show that there is variation in stress-responsive behaviour in a population of wildtype zebrafish. Resilient/susceptible archetypes can be identified at early stages of development, and the attributes are independent of the strain or type of stressful challenge.

### Resilience is a stable and heritable developmental trait

Personality traits are thought to be relatively stable over time (*12*). Therefore, we performed a longitudinal study to examine whether resilience and susceptibility identified in early larvae are maintained through life as they matured through adulthood (Figure 2A). Firstly, we differentiated larvae into unstressed control, resilient and susceptible groups at 6 dpf and allowed them to recover. At 7 dpf, we subjected the larvae to a physical handling challenge to analyze whether the groups show differences in their response to a second unrelated stressor as compared to unstressed controls (Figure 2A, 2B). While the control and susceptible groups showed a decrease in locomotive behaviour when stressed, the resilient larvae did not show a significant reduction (Figure 2B; *p<0.0065*, n=12, 2-way ANOVA; *p<0.0001* control unstressed vs. stressed, susceptible unstressed vs stressed, Sidak’s post-hoc multiple comparisons test).

**Figure 2:**
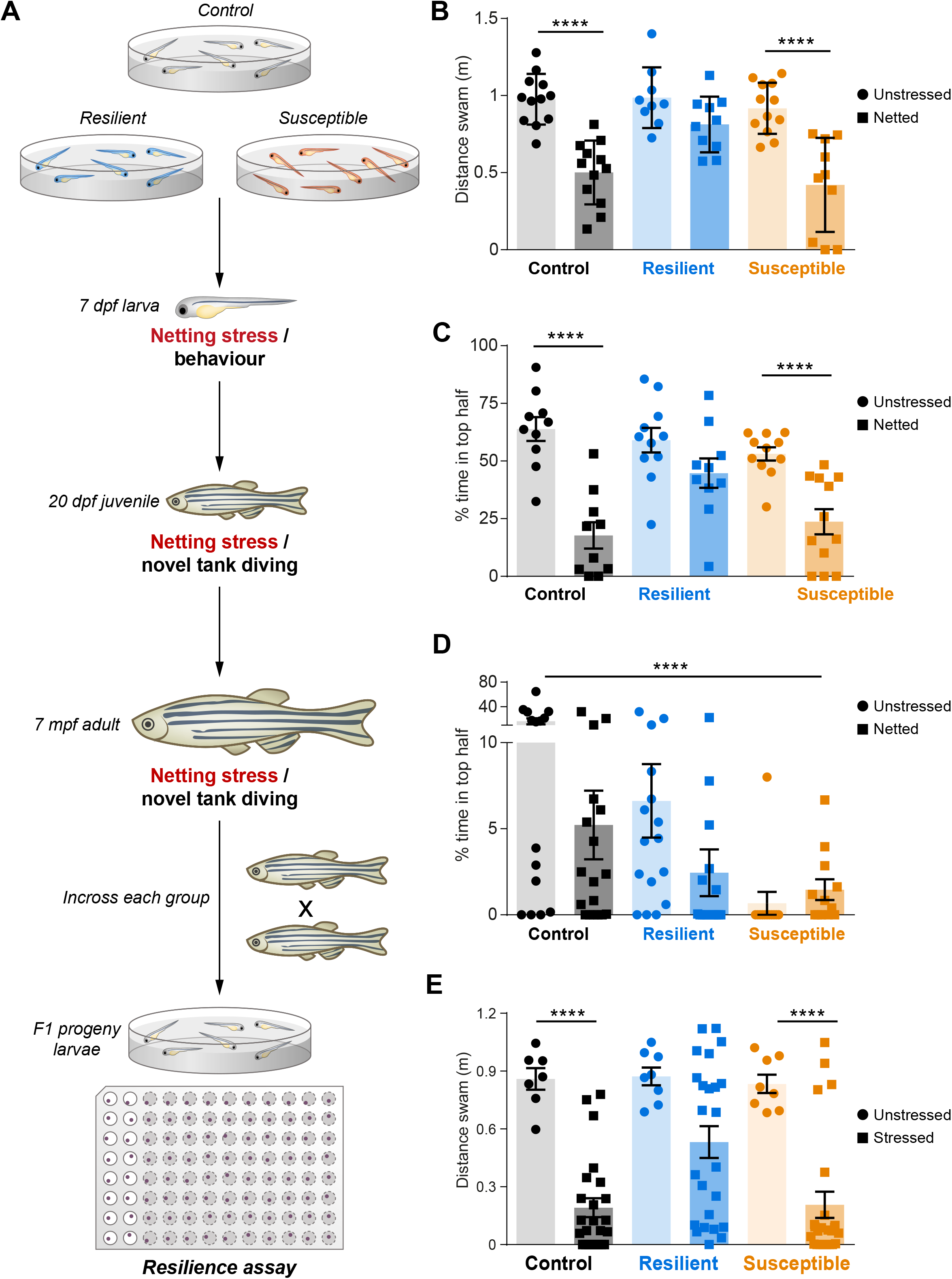
Resilience/susceptibility are persistent and heritable traits. (**A**) Scheme detailing longitudinal persistence and assessment of heritability of traits. Larvae that were separated into control, resilient and susceptible groups at 6 dpf were tested again at 7 dpf, 20 dpf and 7 mpf to analyze the persistence of the attributes. Each group was incrossed and the F1 progeny were tested for heritability. At each stage, every group first was divided into an unstressed sub-group and another subgroup which was removed from water (netted) just before the task. This was followed by behavioural analysis under mildly stressful conditions (novel environment) in accordance with the two-step behavioural paradigm, to analyze the effect of the major stress on behaviour. (**B**) Control, resilient and susceptible groups were subjected to netting stress at 7 dpf, where part of each group was removed out of water for five minutes (netting), followed by analyzing their recovery behaviour in a 96-well plate in the dark. While control and susceptible groups show a decrease in locomotive activity when stressed, the resilient group does not show any difference upon stress. (**C, D**) Control, resilient and susceptible larvae were grown to juvenile (C) and adult (D) stages, and half of the group was subjected to netting stress for two (C) or one (D) minutes followed by analyzing their behaviour in a novel tank diving task. The control and susceptible juvenile groups showed a stress-dependent drop in exploration, compared to the resilient fish (C). At adulthood, the susceptible group showed the least tendency to explore the top half of the tank (D). (**E**) After reaching sexual maturation, resilient and susceptible fish were incrossed with their respective archetype and their F1 progeny were tested for trait heritability. Locomotive behaviour of F1 progeny of the respective archetype were tested either following 20 minutes osmotic stressful challenge or under naïve ‘unstressed’ condition. Of the three groups, the progeny of the control and susceptible fish showed a stress-dependent drop in activity, while the progeny of the resilient fish displayed superior recovery from stress. Data presented as mean±SEM, **** p<0.0001, ***p<0.001, **p<0.01

To further test the stability of these traits, we grew the same larvae under comparable conditions and subjected them to the novel tank diving task as juveniles and adults (Figure 2A, 2C, 2D). This task is widely used to study anxiety-like stress responsive behaviour following introduction of zebrafish to a novel environment (*13, 14*). Typically, when introduced into a tank, the fish dive to the bottom, and thereafter slowly begin to explore the rest of the tank as they acclimate. Stressed fish tend to spend more time in the bottom of the tank. We analyzed the response to novel environment of both, naïve fish that were introduced directly into the tank (unstressed) as well as pre-stressed fish subjected to two minutes netting prior to the test. When examining 20 dpf juvenile fish of control and susceptible groups, we found that netting stressor induced a marked decrease in the percentage of time the fish spent in the top half of the tank. In contrast, the percentage of time the resilient fish spent in the top half was unaffected by the stressful challenge indicating a superior stress coping ability (Figure 2C; *p=0.0054*, n=12, 2-way ANOVA; *p<0.0001* control unstressed vs. stressed, *p<0.001* susceptible unstressed vs. stressed, Sidak’s post-hoc multiple comparisons test).

Testing of resilient and susceptibile fish later in life, revealed that the observed traits remained stable in sexually matured seven months-old adult fish. Adult susceptible fish spent very little time exploring the top half of the tank indicating higher anxiety than the control or resilient fish (Figure 2D, *p<0.0085*, n=18, 2-way ANOVA; *p<0.001* control unstressed vs. stressed, Sidak’s post-hoc multiple comparisons test). Notably, the resilient group had higher proportion of males and the susceptible group had more females (Figure S2A; *p=0.4796* control, *p=0.0009* resilient, *p=0.3785* susceptible; N=6, n∼30 per group per experiment, Chi-square test). Hence, though gender was not specified at the time we tested the larvae at 6 dpf, the archetype of an individual which undergoes early life stress could be a contributing factor in sex determination.

Next, we examined whether resilience and susceptibility are heritable traits. To this end, we in-crossed each group of fish and analyzed their progeny (Figure 2A). Notably, we found that the embryos from a cross of susceptible fish showed inferior quality with more than half the embryos dying within the first 48 hours (Figure S2B; *p<0.0001* control vs. resilient and susceptible; N=3, n∼300 per group per experiment, two-tailed Z test). However, we observed no obvious morphological differences between the surviving F1 progenies of susceptible and resilient parents. When we subjected F1 progenies to osmotic stress and monitored their behaviour, the control and susceptible groups showed an expected reduction in activity, while the resilient larvae were not significantly affected by the stress (Figure 2E, *p=0.0024*, n=24, 2-way ANOVA; *p<0.01* control and susceptible unstressed vs. stressed, Sidak’s post-hoc multiple comparisons test). Following quantification the percentage resilience across multiple experiments, we found that the F1 progeny of the resilient fish consistently showed higher resilience in the population (Figure S2C).

Taken together, we show that resilience and susceptibility, identified in 6 dpf wildtype zebrafish larvae, are stable and heritable traits.

### Mutations in mammalian resilience-associated factors impairs resilience of zebrafish larvae

To examine whether our resilience paradigm displays characteristics of resilience observed in rodents and humans, we tested the transcriptional response of resilient and susceptible larvae focusing on factors associated with resilience and/or stress-response in mammals. We firstly categorized 6 dpf larvae as resilient or susceptible, let them recover for a day and thereafter subjected them to an acute stressor and extracted RNA from whole larvae 30 minutes after the stressful challenge. Of the factors tested, we observed significant differences in the expression of *neuropeptide Y (npy)* (*p=0.0008*, n=3, one-way ANOVA), the transcriptional repressor *rest* (*p=0.0092*, n=3, one-way ANOVA), and the neuropeptides, oxytocin (*oxt)* (*p=0.0241*, n=3, one-way ANOVA) and vasopressin (*avp)* (*p=0.0186*, n=3, one-way ANOVA) (Figure 3A). Stress-induced expression of these factors was lower in the susceptible group in comparison to the resilient group. This shows that differences in the expression of genes associated with stress response in resilient and susceptible larvae are analagous to molecular changes observed in mammals.

**Figure 3:**
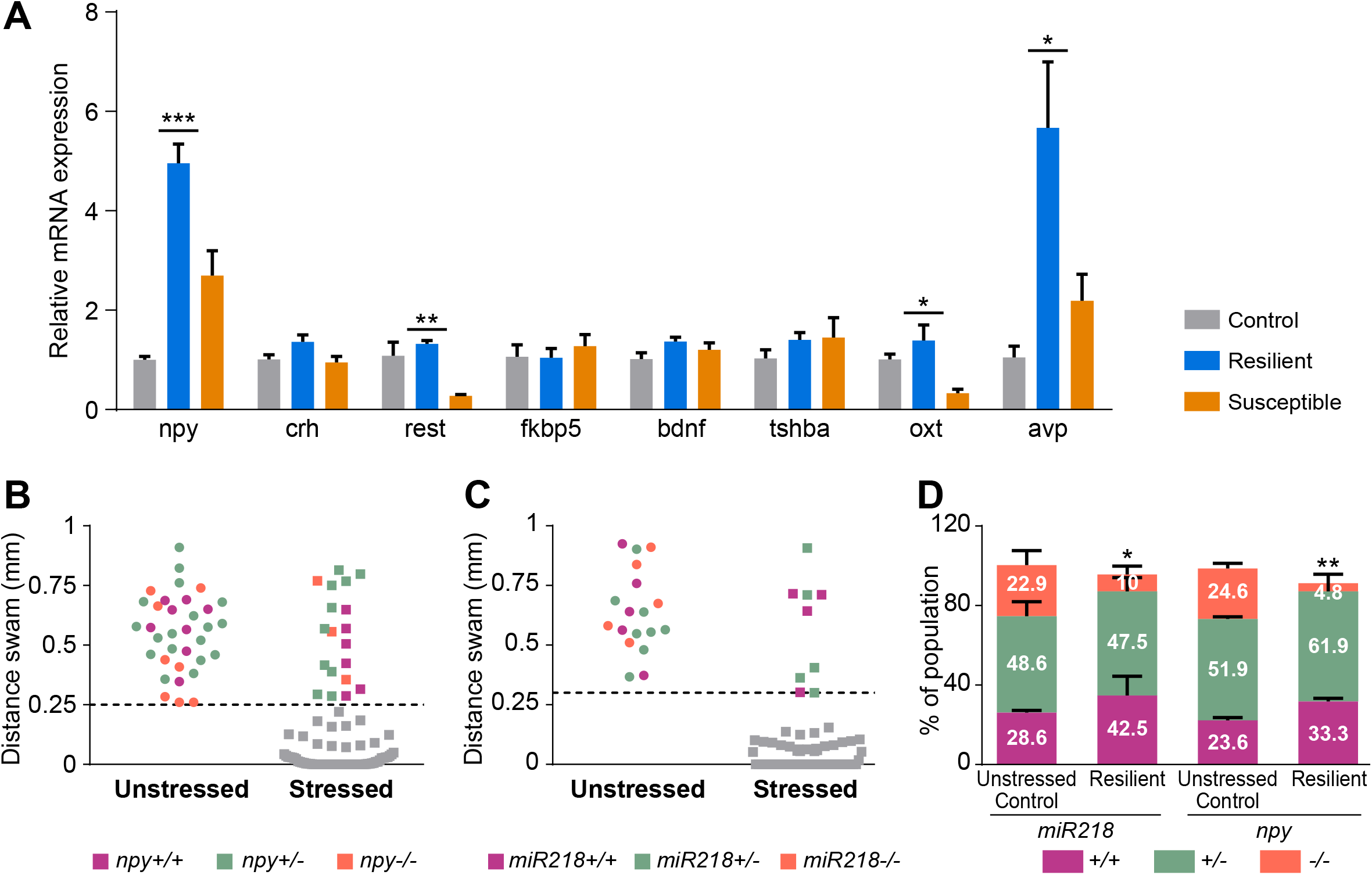
Stress resilience is modulated by neuropeptides. (**A**) Gene expression analysis of factors known to be associated with stress resilience and stress response. Six dpf larvae were differentiated into control (unstressed), resilient and susceptible groups as described in Figure 1. After 24 hours of recovery, the three groups were subjected to 5 minutes osmotic stress and stress-responsive gene expression was analysed after 30 minutes. (**B-D**) Resilience of mutants lacking functional NPY and miR218. Heterozygous mutant fish were crossed and the progeny were subjected to the resilience assay. Unstressed control and resilient fish were blindly genotyped after the assay to analyze the actual proportion of wildtype: heterozygotes: homozygotes. (**B, C**) Dot plots representing the distance swam during the recovery phase of the progeny of heterozygous *npy* (B) and *miR218* (C) mutant larvae that were left unstressed or pre-subjected to a major stressor. Genotyping of unstressed control and resilient larvae showed presence of all genotypes in the control group, but lesser than the expected 25% enrichment of *npy-/-* (B) and *miR218-/-* (C) larvae in the resilient population. (**D**) Data from multiple experiments showing proportions of wildtype:heterozygotes:homozygotes genotypes in unstressed control vs. resilient progenies derived from *npy* and *miR218* heterozygotes was analyzed using Chi-square test. The unstressed group had the expected 1:2:1 ratio, while the *npy* and *miR218* homozygous mutants were significantly underrepresented in the resilient population. Data presented as mean±SEM *p<0.05, **p<0.01, ***p<0.001

We next examined whether larval resilience is affected by loss-of-function of the resilience-associated factors NPY and miR218 that have been positively correlated with stress resilience (in rodents and human) and whose low level in the forebrain promote susceptibility and depression (*5, 15*). As positive effectors of resilience, larvae harbouring genetic deficiencies in *npy* and *miR218* would be less resilient, and hence, underrepresented in the resilient group. To examine this, we tested the progeny of heterozygous mutant *npy* (*16*) and *miR218* (Figure S3A-D; *p=0.04*, one-way ANOVA for gene expression analysis) in order that full siblings were compared to each other in our resilience assay simultaneously. Following this, we genotyped the unstressed control and resilient populations and compared the ratios of wildtype:heterozygote:homozygote larvae to the expected Mendelian ratio of 1:2:1, respectively using a Chi-square test. This analysis showed that the proportion of genotypes in the unstressed group was according to an expected Mendelian ratio (Figure 3D; *p=0.8794 miR218* control, *p=0.9615 npy* control; N=3, n=40, Chi-square test). In contrast, both *npy-/-* and *miR218-/-* mutants were significantly underrepresented in the resilient population suggesting that NPY and miR218 positively affect zebrafish resilience (Figures 3B-D; *p=0.0139 miR218* resilient, *p=0.0029 npy* resilient; N=3, n=40, Chi-square test). These experiments confirmed that *npy* and *miR218* impact stress resilience in zebrafish larvae, similar to their previously suggested roles in mammals.

### Resilient larvae display unique stress-responsive transcriptional response

Having established that mammalian resilience-associated genes play a similar role in zebrafish, we sought to take an unbiased approach to identify additional factors contributing to resilience. We differentiated larvae into resilient and susceptible groups at 6 dpf and on the following day, larvae were collected either at basal conditions or 30 minutes following an acute stressful challenge, which we previously showed to evoke a peak in the transcription of *crh* (*11*) (Figure 4A). Bulk mRNA sequencing of the four groups of larvae showed that resilient fish display a unique stress-responsive gene expression changes (Figure 4B; n=5 per group). Out of the approximately 25000 genes expressed in larvae at this stage, approximately 250 were downregulated and 100 were upregulated in the resilient stressed larvae in comparison to the three other groups (**Supplementary table S1**; fold change > 1.5; adjusted p value < 0.05). Our finding that resilient larvae show specific stress-responsive transcriptional changes correlates well with previous studies showing that resilience is an active process involving more transcriptional changes than susceptibility (*17*).

**Figure 4:**
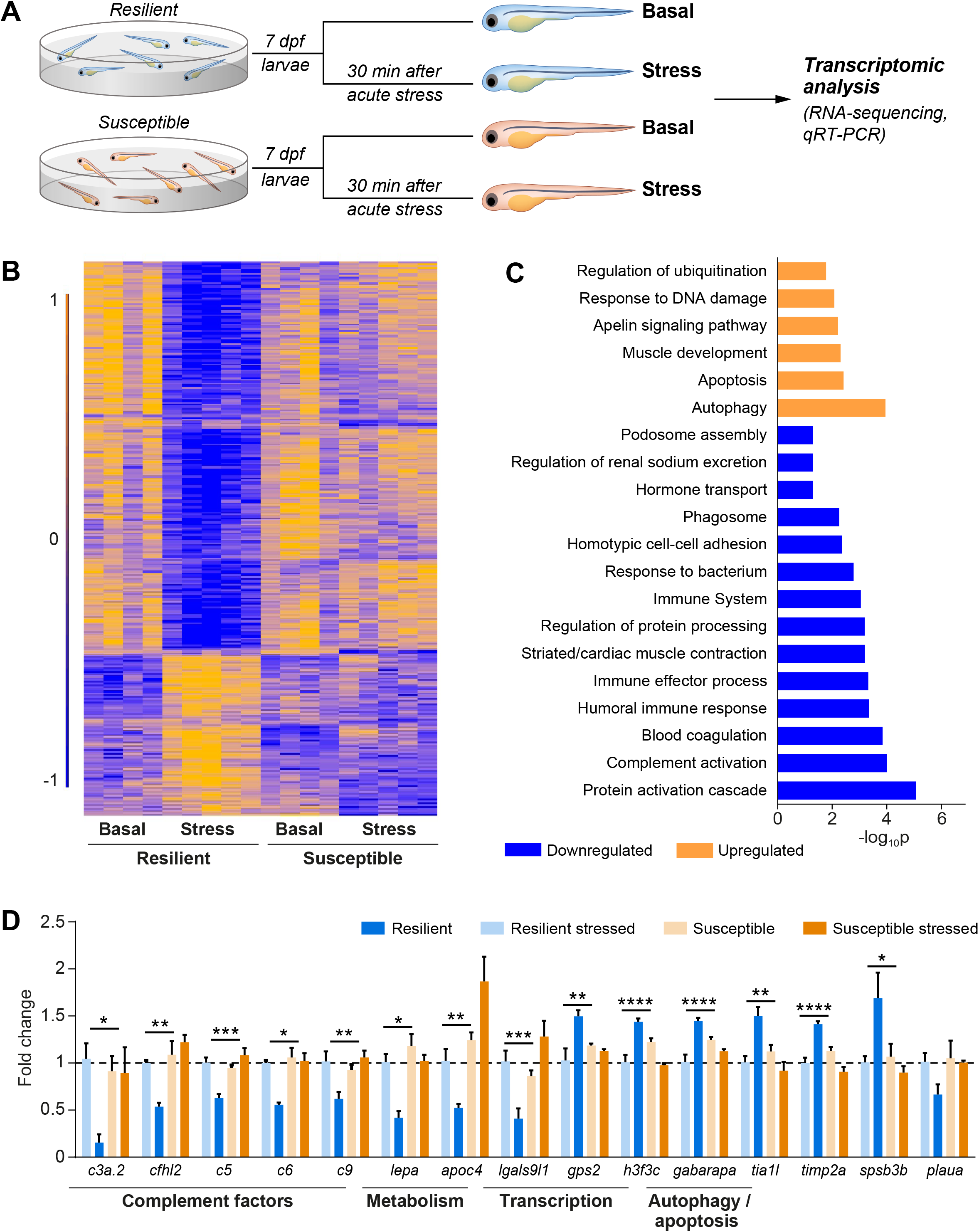
Resilient larvae show unique stress-responsive changes in gene expression. (**A**) Scheme showing the experimental design. Larvae were differentiated into resilient and susceptible groups at 6 dpf followed by 24 hours recovery. At 7 dpf, each archetype was either subjected to 5 minutes osmotic stress (*Stressed*) or left unstressed (*Basal*) and stress-responsive gene expression changes in both groups were analyzed by mRNA sequencing 30 minutes thereafter. (**B**) Heatmap of differentially expressed genes in resilient and susceptible individuals comparing basal and stressed states. Each row indicates expression of one gene relative to the other larvae (dark blue, −1, dark yellow, 1, scale is in standard deviation units for each gene). (**C**) Functional enrichment analysis showing gene ontology terms and pathways. Terms associated with significantly up-(yellow) and downregulated genes (blue) were determined using Metascape and Enrichr. GO terms related to the immune system, and specifically related to complement activation were enriched for the downregulated genes in the stressed resilient larvae (highlighted in red). (**D**) Quantitative RT-PCR validation of relevant genes that were up-regulated or down-regulated in the resilient larvae following stress. Multiple components of the complement pathway were checked, and all the factors tested show a significant downregulation in resilient larvae when stressed. mRNA levels were normalized to beta-actin levels and are represented as fold inductions, with the value obtained with the resilient basal larvae set at 1. Data presented as mean±SEM, **** p<0.0001, ***p<0.001, **p<0.01, *p<0.05

Functional enrichment analysis of differentially expressed genes revealed that, when stressed, resilient larvae downregulated genetic factors involved in immune response and renal function, and upregulated factors associated with autophagy, apoptosis, ubiquitination and DNA damage response (Figure 4C, 4D**, Supplementary table S2**). Upon a closer look, we observed that almost all members of the complement pathway were downregulated (Figures 4C, S4A **and** 4D). Representative genes that were up- or downregulated in the RNA-sequencing experiment were further validated by qRT-PCR (Figure 4D; *p=0.047 c3a.2, p=0.005 cfhl2, p=0.0005 c5, p=0.0103 c6, p=0.0062 c9, p=0.033 lepa, p=0.0032 apoc4, p=0.001 lgals9l1, p=0.0036 gps2, 0<0.0001 h3f3c, p<0.0001 gabarapa, p=0.0012 tia1l, p<0.0001 timp2a, p=0.0199 spsb3b, p=0.2317 plaua* n=4, two-way ANOVA). The above gene expression changes in both the resilient and susceptible larvae was independent of the type of stressor, as netting challenge induced the same gene expression response as the osmotic stress (Figure S4B; *p=0.0017 lepa, p=0.0034 c3a.2, p=0.0073 cfhl2, p=0.0189 lgals9l1, p=0.0511 gps2, p=0.0099 gabarapa, p=0.0077 tia1l, 0.0243 h3f3c, p=0.9243 spsb3b*, n=4, unpaired t-test). Hence, irrespective of the nature of the stressor, resilient larvae display a specific transcriptional response, presumably to actively cope with the effects of stress.

Since our transcriptome analysis was performed on mRNA extracted from whole larvae, we used an existing zebrafish larval single-cell RNA sequencing (scRNA-seq) atlas to determine the tissue specificity of selected candidate genes showing differential expression in resilient vs. susceptible larvae (*18*). For this analysis, the expression of each of the significantly up- and downregulated genes revealed herein, was visualized on the dimensional reduction plot of the published 5 dpf scRNA-seq atlas (*18*) and the genes were assigned to specific cell clusters. This analysis revealed that genes that were downregulated in the resilient fish following stress were conspicuously highly expressed in hepatocytes and monocytes/macrophage cell types (Figure 5A**, Supplementary table S3**). In particular, members of the complement pathway, which were downregulated in the resilient larvae following stress, displayed liver-specific expression (Figure 5B, 5C**, Supplementary table S3**).

**Figure 5:**
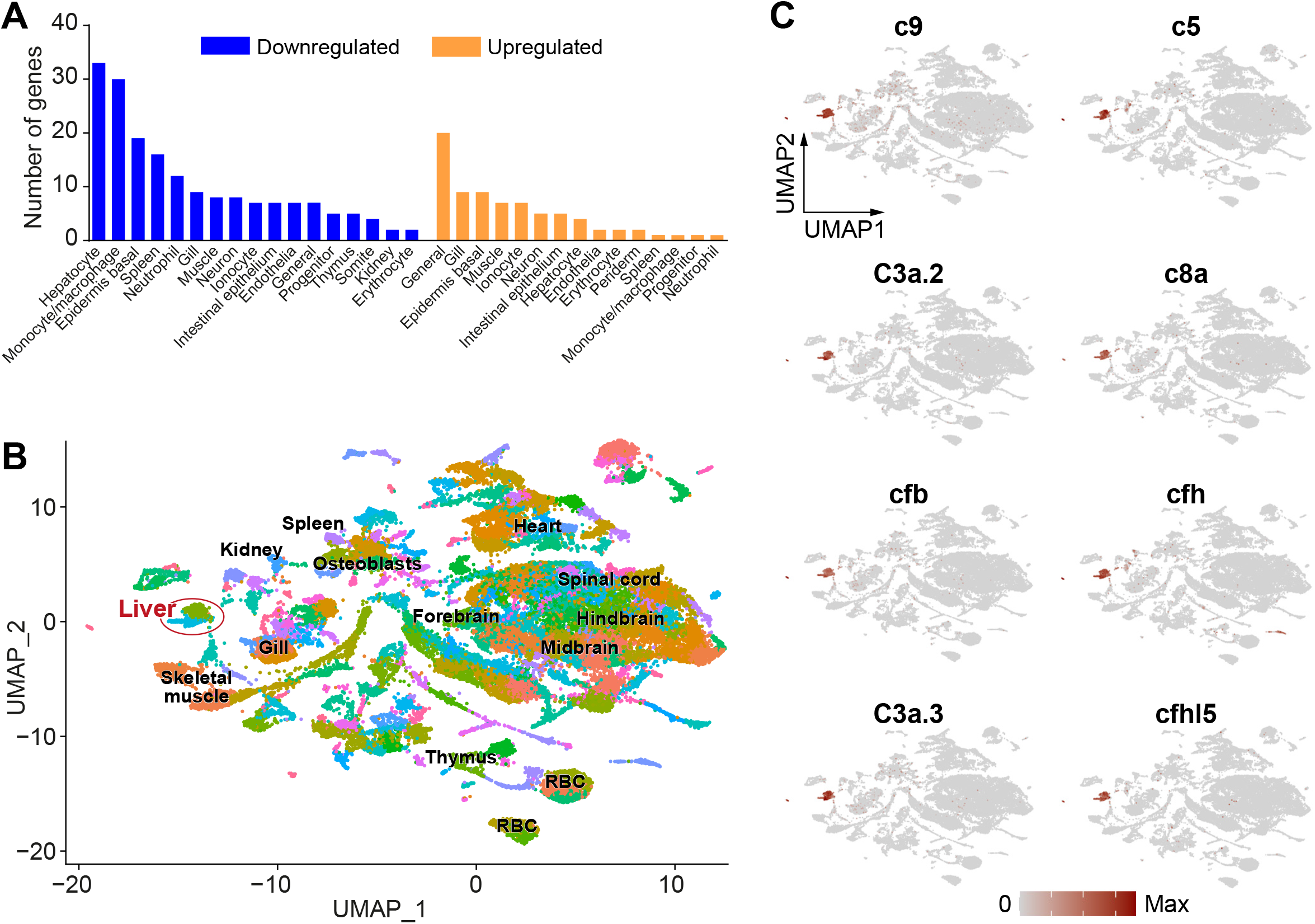
Complement factors downregulated in stressed resilient larvae are expressed in the liver. (**A**) Tissue specificity revealed by visualization of significantly down- and upregulated genes on a dimensional reduction plot of a published scRNA-seq atlas (*18*). The histogram shows the number of upregulated and downregulated genes in each tissue and/or cell types. (**B**) Uniform manifold approximation and projection (UMAP) showing topological distribution of single cell gene expression clusters of whole 6 dpf larvae. The UMAP was reconstructed from a published scRNA-seq atlas (*18*). The different colors represent different clusters assigned to cell types and major representative tissues are indicated. (**C**) Feature plots in which the expression of selected complement factors which were downregulated in resilient larvae vs. susceptible larvae is highlighted in the UMAP plot showing distinct expression of these factors in liver hepatocytes.

Taken together, our gene expression analysis of resilient and susceptible archetypes revealed unique tissue-specific stress-responsive transcriptional changes in both, central and peripheral organs.

### Perturbations of the complement pathway affect resilience

As mentioned above, we observed that the innate immune complement pathway showed significant enrichment and multiple members of this pathway were downregulated in resilient larvae subjected to stress. The complement system is made up of a large number of distinct plasma proteins that react with one another to opsonize pathogens and induce a series of inflammatory responses that help to fight infection (*19*). As shown above, multiple complement factors are synthesized in the liver and are known to be secreted to general circulation. These factors have been associated with systemic physiological stress responses and used as biomarkers for anxiety and depression (*20–22*). Moreover, peripheral activation of innate immunity by lipopolysaccharide in animal models is known to induce “sickness-behaviour”, which is associated with abnormal stress response, including depression (*23*). Hence, we hypothesized that the complement pathway is a critical determinant in the establishment of resilience.

Since the expression of complement factors was downregulated in the resilient larvae, we generated mutants which lacked the crucial complement factors, *c5* and *c9* (Figures S5A-G; *p<0.0001* one-way ANOVA for gene expression analysis of *c5* and *c9*). We anticipated that both c5 and c9 mutants would be enriched in the resilient population. To examine this, we subjected the progenies of heterozygous fish to our stress resilience assay and compared the ratios of wildtype:heterozygote:homozygote larvae in both the unstressed and resilient populations to the expected Mendelian ratios using a Chi-square test (Figures 6A and 6C). The expected Mendelian genotypes ratio of 1:2:1 [(+/+);(+/-);(-/-)] was found in the unstressed population, while the resilient population had significantly greater percentage of mutants across multiple experiments (Figures 6B and 6D; *p=0.2351 c5* control, *p=0.014 c5* resilient, *p=0.7941 c9* control, *p=0.0021 c9* resilient; N=9, n∼70 control/115 resilient, Chi-square test). These experiments indicate that stress-induced suppression of the complement pathway is a key positive determinant of early life stress resilience.

**Figure 6:**
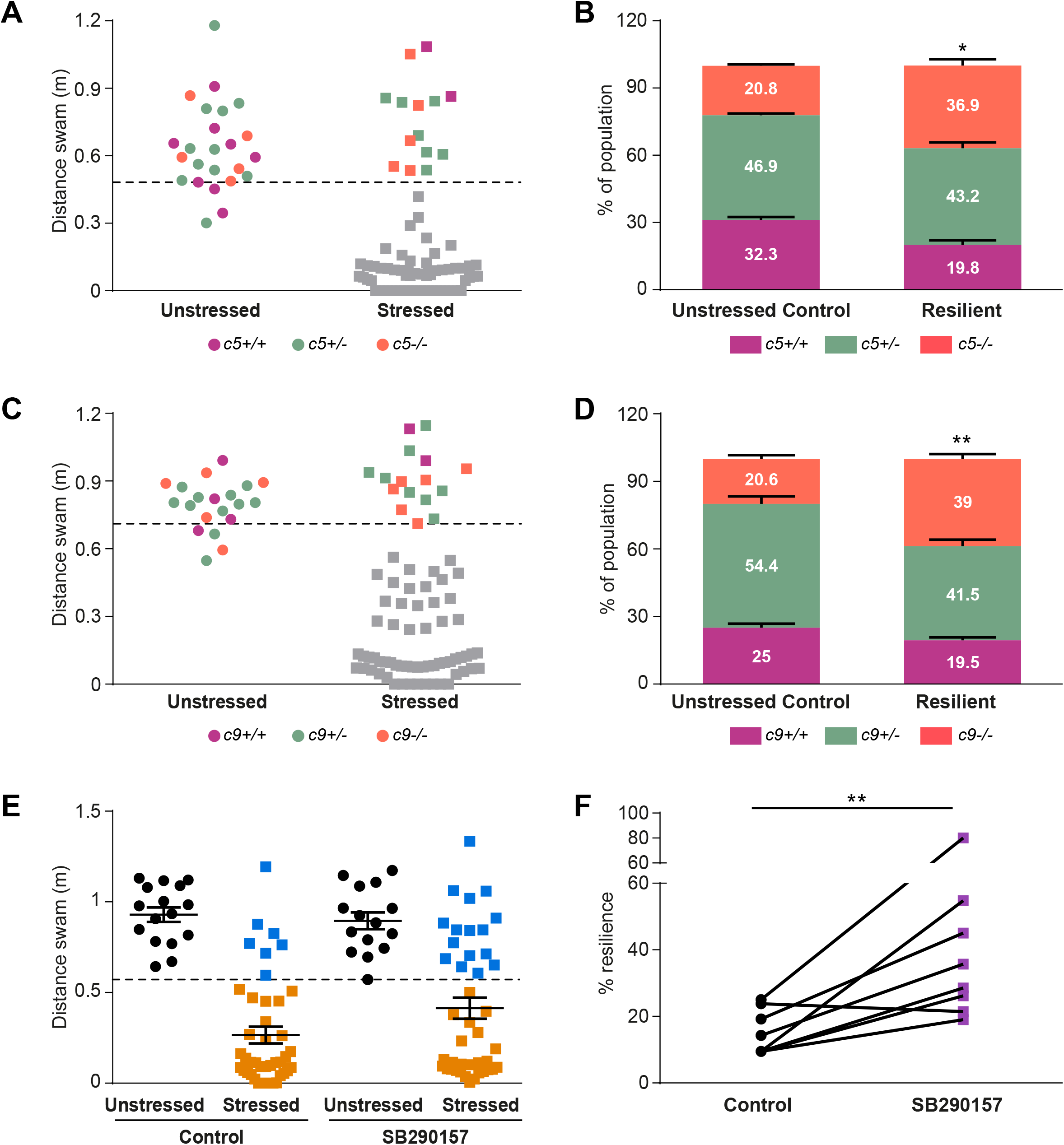
Complement inhibition is a key determinant of early life stress resilience. (**A-D**) Heterozygous *c5* and *c9* mutants were crossed, the progeny divided into unstressed control and pre-stressed, and subjected to the resilience assay. Unstressed and resilient larvae were genotyped after the assay and the actual proportion of wildtype:heterozygotes:homozygotes in comparison to the expected ration of 1:2:1 was analyzed using a Chi-square test. *c5-/-* (**A,B**) and *c9-/-* (**C,D**) larvae are larvae show greater than the expected 25% representation in the resilient population. (**E, F**) Larvae were treated with SB290157, a C3aR antagonist for 12 hours and the effect on resilience was tested at 6 dpf. Control and SB290157 treated larvae were either stressed or left unstressed and subjected to the resilience paradigm (Figure 1). (**E**) In comparison to the control group, the treated group showed greater number of resilient fish (blue squares-compare control and SB290157 groups). (**F**) Treatment with SB290157 showed a consistent increase in the percentage of resilience over multiple experiments. Each dot represents one experiment. Data presented as mean±SEM, **p<0.01, *p<0.05

To further substantiate this finding, we analyzed the effect of pharmacological inhibition of the pathway on the percentage resilience in a population. We noticed that c3a was the most robustly downregulated complement factor in the resilient archetype (Figures 4D and S4A) and hence, treated larvae with SB290157, a specific C3aR complement receptor antagonist (*24*). Overnight treatment with SB290157 led to a robust increase in the number of resilient larvae in the treated population across multiple cohorts (Figure 6E and 6F; *p=0.0084*, N=8, n=6 unstressed/42 stressed per experiment, paired t-test). When we treated with rosmarinic acid, an inhibitor of C3 and C5 convertase (*25, 26*), we observed a similar increase in resilience (Figure S6; *p=0.0022*, N=8, n=6 unstressed/42 stressed per experiment, paired t-test).

Taken together, we present a new method to quantify stress resilience in young zebrafish larvae and subsequently classify them as being resilient or susceptible to stress. We further show that these traits are stable throughout life and independent of the type of stressor. Resilience is augmented by NPY and miR218 and attenuated by innate immune complement factors.

## Discussion

All individuals are facing different stressful situations, and the ability to effectively rebound from stress varies across a given population. An immense body of previous research shows that early life experiences impact stress-responsive behaviour later in life (*6, 27, 28*). However, most studies aimed at understanding the basis of resilience/susceptibility to date have been performed on adult mammals because it is challenging to analyze behaviour in young mammals whose development occurs *in utero* and shortly after birth. Hence, whether young vertebrates already display attributes of resilience and susceptibility is unknown. Zebrafish embryos develop from eggs fertilized externally and these fish show robust quantifiable stress-responsive behavior as early as 6 days post fertilization (dpf). Using zebrafish as a model, we have designed and established a high-throughput assay to identify stress resilience/susceptibility early in life.

Our study shows that resilience and susceptibility can be measured as stable traits from early stages of development in vertebrates. We employed a stringent statistical method to reliably identify resilient individuals and using this method we observe that 10-20% of individuals in a wildtype population show stress resilient behaviour, which is comparable to the reported ∼35% of resilient mice following chronic social defeat stress paradigm (*29, 30*).

Previous research has shown that children of individuals with psychological illnesses have greater tendency towards stress susceptibility (*31*). However, young mammals are extremely dependent on parental care during their early life stages and therefore, it is difficult to distinguish between the contribution of parental care to susceptible tendencies from the effect of genetics. Our assay could be useful for further research on the contribution of genetic factors to stress resilience without the interfering confound of parental care as we show that resilience/susceptibility are inherited traits in zebrafish, whose embryos, larvae and juveniles develop and grow independently of parental care.

Mechanistically, we demonstrate that NPY and miR218 play a role in determining early life resilience. NPY and miR218 have been previously associated with resilience in mammals (*2*). Thus, the levels of NPY and miR218 are positively correlated with stress resilience and NPY attenuates depression-like behaviours in rodent models and humans (*5, 15, 32, 33*). Here, we show that the role of these factors is conserved across species and that they begin to function very early in life to establish stress resilience.

In addition to specific factors and brain areas regulating stress resilience centrally, many peripheral factors like the immune system (both innate and adaptive) and gut microbiota have been shown to influence resilience/susceptibility to stress (*2, 34*). Of the innate immune factors, inflammatory cytokines are directly correlated with depression; anti-inflammatory therapies elicit antidepressant effects and vice versa (*35–37*). Stress activates inflammatory signaling pathways probably through epigenetic mechanisms (*38–40*). Our unbiased transcriptome approach from individual larvae classified as resilient or susceptible, revealed a specific and previously unknown link of complement activation with stress resilience. The expression of multiple components of the innate immune complement system was downregulated in resilient individuals in response to stressful challenge. We further showed that pharmacological inhibition of C3 action as well as knocking out C5 and C9 genes improved resilience. While dysfunctional complement activation has been correlated with multiple psychological conditions, such as depression and anxiety (*20, 41, 42*), our results show that fine control over the activity of critical components of the complement cascade during early-life underlies resilience.

The underlying mechanism by which complement factors affect resilience is yet to be determined. While normal functioning of the complement pathway is critical for combating infections, overactivation can lead to tissue damage by means of the membrane attack protein complex (*43*). Thus, mitigation of complement factors following stressful challenges may prevent unwanted tissue damage, and hence greater resilience. An alternative possible mode of action could be independent of the classical membrane attack complex but rather mediated by the binding of circulating complement factors to tissue localized complement receptors. For instance, C3a complement receptor is expressed in the pituitary and its activation by the cognate complement ligands stimulates the hypothalamo-pituitary-adrenal stress axis and consequently, the release of adrenocorticotropin hormone (ACTH) (*44*).

It is increasingly being recognized that stress influences the immune system and peripheral immune cells impact brain circuits making a strong case for neuroimmune interactions regulating behaviour. Hence, an important issue to be addressed in future studies is the site of action of complement factors in the context of stress resilience. Many of these factors like C8, C9, CFB are synthesized in the liver and thereafter secreted to the circulation (*18, 45, 46*). Hence, the effect of liver-specific complement factors on resilience is likely due to their peripheral function. On the contrary, the effect of complement downregulation on resilience could be a direct consequence of tissue-specific events. For instance, specific depletion of components of the pathway in brain regions or cells affects synaptic refinement and behavior (*47, 48*). Hence, the site and mechanism by which complement downregulation leads to increased stress resilience remains to be studied further. Finally, further studies are required to understand if there exists a crosstalk between the regulation of stress resilience by central neuropeptides, such as NPY and peripheral immune factors.

To conclude, we report that resilient and susceptible traits can be identified early in life as stable traits. These traits are regulated by a combination of neuropeptides and miRNAs that function centrally and by innate immune complement factors that function peripherally.

## Materials and methods

### Zebrafish husbandry and maintenance

Zebrafish were raised and bred according to standard protocols. Husbandry and experimental procedures were approved by the Institutional Animal Care and Use Committee of the Weizmann Institute, Israel (#01420120-2 and # 09570119-3). Adult fish were maintained in mixed sex groups in a Techniplast system at 28 C, pH 7 and conductivity 1000 µS/cm. Embryos were grown at 28 C in 0.3X Danieau’s medium at constant uncrowded densities (17.4 mM NaCl, 0.21 mM KCl, 0.12 mM MgSO4, 0.18 mM Ca(NO3)2, 1.5 mM HEPES, pH 7.4). Larvae, juvenile (3-8 weeks old) and adult zebrafish (3-6 months old) of both sexes were used in this work.

### CRISPR mutagenesis

To generate mutants of miR218a-1 (ZFIN ID, ZDB-ALT-190814-7) and complement factors c5 and c9 (ZFIN ID, ZDB-ALT-220120-2 and ZDB-ALT-220120-3), CRISPR guide RNAs were designed using Benchling, CRISPOR and CRISPRDirect. Two gRNAs were designed for each gene that was targeted, one upstream of the translation start site and one after the last possible alternative translation start site (*49*) (sequence-supplementary table). gRNAs were procured from Integrated DNA Technologies and Cas9 protein was made by the Weizmann Institute’s Protein purification unit by expressing from the pET-28b-Cas9-His plasmid (*50*). A mixture of 1.5 µM gRNA and 250 µg Cas9 protein was incubated at 37 C for 10 minutes for complex formation and injected into AB embryos at the one-cell stage. Injected fish were raised to adulthood and crossed. The progeny were screened for presence of large deletions resulting from cleavage at both CRISPR target sites and germline transmission. The exact nature of the mutation was identified by Sanger sequencing. Progeny from founders outcrossed for maintenance and further experiments were performed with subsequent generations.

### DNA extraction and genotyping

Genomic DNA was extracted from tail fin clips or 24 hpf embryos by lysis in 50 mM sodium hydroxide, incubation for 30 minutes at 95 C and neutralization using 100 mM Tris-HCl pH 8. gDNA prepared thus was used for amplification of the region of interest by PCR (primer sequences-supplementary table). PCR products were separated on a 1% agarose gel by electrophoresis and visualized using a UV transilluminator. If required, the DNA was extracted from the agarose gel using a gel extraction kit (Hylabs, Israel) and sequenced by the Sequencing Unit at the Weizmann Institute of Science.

### Larvae stress resilience assay

Experiments were performed on 6 dpf wildtype AB larvae (unless otherwise mentioned) between 8:30 AM and 3:00 PM. Larvae were allowed to acclimate to the test room for 20 minutes before the task. For osmotic stress, larvae were treated with 50% artificial sea water (3.5 g Instant Ocean sea salt in 200 mL Danieau’s buffer) for 20 minutes unless stated otherwise. Unstressed control larvae were handled similarly, but treated with Danieau’s buffer. Following washes, the larvae were transferred into individual wells of a 96-well plate, with each well containing 0.3 ml of Danieau’s buffer. Recovery of the larvae was recorded for 1 hour in the Noldus Daniovision system and analyzed using Ethovision XT14.0 (Noldus, Netherlands). The lag between the stress and beginning of the recording was kept as minimal as possible. Distance swam was binned in 5 minute intervals and data was exported for further statistical analysis. For netting stress, larvae were transferred to a cell strainer and removed from water for a period of five minutes followed by handling and analysis comparable to the osmotic stress assay.

For analysis of heritability, the unstressed controls, resilient and susceptible groups were raised to adulthood and each group was incrossed. Six dpf progeny from each group were divided into two sub-groups each: unstressed and stressed. The stressed groups were subjected to the osmotic stress assay and stress recovery behaviour was recorded and analyzed.

### Pharmacological treatments

Rosmarinic acid (10 µM, 12 hours) and SB290157 (10 µM, 12 hours) treatments were performed to analyze the effect of inhibition of the complement pathway on recovery from stress. Following treatments, the control and treated larvae were divided into two sub-groups each: unstressed and stressed. The stressed group was subjected to the osmotic stress assay described above and stress recovery behaviour was recorded and analyzed.

### Novel tank diving assay

Juvenile (20 dpf) or adult zebrafish (7 months old) were habituated to the test room for 1 hour before the task. Anxiety-like behaviour was measured using the novel tank diving assay as described earlier (*11*). The assay was performed in an enclosure with lighting from the top and insulated from any audio or visual inputs. Videos were acquired using a 2M360-CL camera and recorded with Streams5 software (IO industries, Ontario). Videos were analyzed using Ethovision XT14 (Noldus, Netherlands).

### Transcriptomic analyses

To perform gene expression studies, larvae were differentiated into resilient and susceptible groups at 6 dpf using the above-mentioned assay and let to recover from the stress overnight. At 7 dpf, the groups were divided into two: basal, which was kept under normal conditions, and stressed, in which larvae were stressed for 5 minutes using 50% artificial seawater. Both groups were collected 30 minutes following the stress. Single larvae were transferred into previously prepared 1.5 ml microcentrifuge tubes, transferred to ice, and the water was quickly replaced with 100 µl Tri reagent (Molecular Research Center Inc). Total RNA was extracted from individual fish larvae using manufacturer’s protocol. Quality of the RNA was assessed using Nanodrop 260/280 and 260/230 ratios and on a TapeStation 2200system (Agilent Technologies).

### RNA-sequencing

RNA-seq libraries were prepared at the Crown Genomics institute of the Nancy and Stephen Grand Israel National Center for Personalized Medicine, Weizmann Institute of Science, using the INCPM-mRNA-seq protocol. Briefly, the polyA fraction (mRNA) was purified from 500 ng of total input RNA followed by fragmentation and the generation of double-stranded cDNA. After Agencourt Ampure XP beads cleanup (Beckman Coulter), end repair, A base addition, adapter ligation and PCR amplification steps were performed. Libraries were quantified by Qubit (Thermo Fisher Scientific) and TapeStation (Agilent). Sequencing was done on a Nextseq instrument (Illumina) using a 75 cycles high output kit, allocating 20M reads per sample (single read sequencing).

### Analysis

Raw fastq files were analyzed using the User-friendly Transcriptome Analysis (UTAP) pipeline developed at the Weizmann Institute of Science (*51*). Reads were mapped to the reference genome *Danio rerio* (danRer11). Briefly, in the pipeline, adapter and low quality base trimming was performed using Cutadapt, followed by mapping to the reference genome, gene annotation, and quantification by counting reads using STAR. Finally, the DESeq2 package was used to normalize and obtain differential gene expression comparing the 4 groups: resilient basal, resilient stressed, susceptible basal and susceptible stressed (Fold change ≥ 1.5; adjusted p value ≤ 0.05; mean normalized read counts ≥ 5). Heatmap was generated using the Partek Genomics suite. Further analysis of the differentially expressed genes for functional enrichment was performed using Metascape (*52*) and Enrichr (*53*).

RNA-sequencing data is available on Gene Expression Omnibus (GEO; www.ncbi.nlm.nih.gov/geo; Accession number: GSE193433).

Visualization of specific genes using Seurat (*54*) FeaturePlot function of zebrafish larva scRNA-seq atlas, reconstructed from Atlas.rds (Seurat v2) downloaded from https://www.adammillerlab.com/resources, data presented in (*18*).

### qRT-PCR

250 ng of RNA was used for reverse transcription to prepare cDNA with Takara PrimeScript RT Master Mix (Cat no. RR036A). cDNA prepared thus was diluted 1:60 and 4 µL of the diluted cDNA was used for each qPCR reaction with SYBR green (Invitrogen). β-actin was used as the reference endogenous gene for normalization. The qPCR reaction was performed in 96 well plates in a StepOne Plus RealTime system. Primer sequences can be found in supplementary table. Relative gene expression was quantified by the ΔΔCt method.

### Statistical analyses

Data are presented as mean ± standard error of mean (SEM). Analyses were performed using PRISM 6.0 (GraphPad Software Inc, San Diego, USA). Normality of the data obtained after stressing zebrafish larvae was tested by the Shapiro-Wilk test. Depending on the outcome, parametric (ANOVA) or non-parametric tests (Kruskal-Wallis test) were used for analysis. Post-hoc tests (Tukey’s, Dunn’s or Sidak’s multiple comparisons) were performed.

For male-female proportion analysis and to compare observed distribution of wildtype:mutant genotypes after behaviour to expected Mendelian distributions, chi-square test was performed. For analysis of proportion of normal:unfertilized eggs, a two-proportion Z test was performed comparing the progeny of the resilient and susceptible independently to the control. For analysis of survival, long-rank (Mantel-Cox) test was performed. For all tests, p<0.05 was considered significant.

#### Resilient/susceptible differentiation

Behaviour of larvae after 20 minutes osmotic stress was analyzed for one hour. In the rare cases where control unstressed larvae displayed general lack of movement they were excluded from the analysis. When multiple experiments were analyzed for normality using the Shapiro-Wilk test, the data never showed normal distribution. Subsequent log transformation of the values showed that the population separates into two and the distribution is binomial.

For unbiased separation of resilient and susceptible archetypes a statistical method of dynamic thresholding was used by combining two tools: Cohen’s d effect size and Fisher’s exact test analyzing contingency tables. In this manner, each data point was considered as the potential threshold. The larvae that were above and below this point were considered resilient and susceptible respectively and the values of Cohen’s d and Fisher’s exact p were calculated as follows.

Cohen’s d = (Mean_above_ + Mean_below_) / √((Std dev_above_)^2^ + (Std dev_below_)^2^)

Considering the following table,

|  |  |
| --- | --- |
| Number of control larvae above threshold (a) | Number of stressed larvae above threshold (b) |
| Number of control larvae below threshold (c) | Number of stressed larvae below threshold (d) |

Fisher’s exact p = ((a+b)! (c+d)! (a+c)! (b+d)!) / (a! b! c! d! (a+b+c+d)!)

The values of the Cohen’s d and Fisher’s exact p were normalized so as to be within the range of 0 to 1. The threshold where the sum of the two values was the maximum and closest to 2 indicating that the groups were most significantly different at this point, was chosen as the actual threshold. All the fish that showed activity above this threshold were considered resilient and the other larvae with activity below the threshold were considered susceptible. Any resilient larvae that showed activity above the threshold only in the first five minutes but never after this period were considered false positives.

## Acknowledgements

We thank Roy Hofi and fish facility personnel; Dana Robbins for running the RNA-sequencing; Dr. David Prober (California Institute of Technology) for sharing the *npy-/-* mutant; Dr. Ron Rotkopf and Dr. Shalev Itzkovitz for helpful discussions on the thresholding method; Genia Brodsky for the figure graphics. G.L. lab is supported by the Israel Science Foundation (#349/21); US-Israel Bi-National Science Foundation (#2017325); Israel Ministry of Science (#3-16548); Hedda, Alberto, and David Milman Baron Center for Research on the Development of Neural Networks; Sagol Institute for Longevity Research; and Maurice and Vivienne Wohl Biology Endowment. G.L. is an incumbent of the Elias Sourasky Professorial Chair.

## Authors’ contribution

A.S and G.L designed the study. A.S designed and performed the experiments and analysed data. M.G performed experiments with the *npy-/-* mutant, S.A generated the *miR218-/-* mutant, N.W contributed to analyzing RNA-sequencing data. A.S and G.L wrote the manuscript.

## Supplementary information

### Supplementary figure legends

**Supplementary figure S1:**
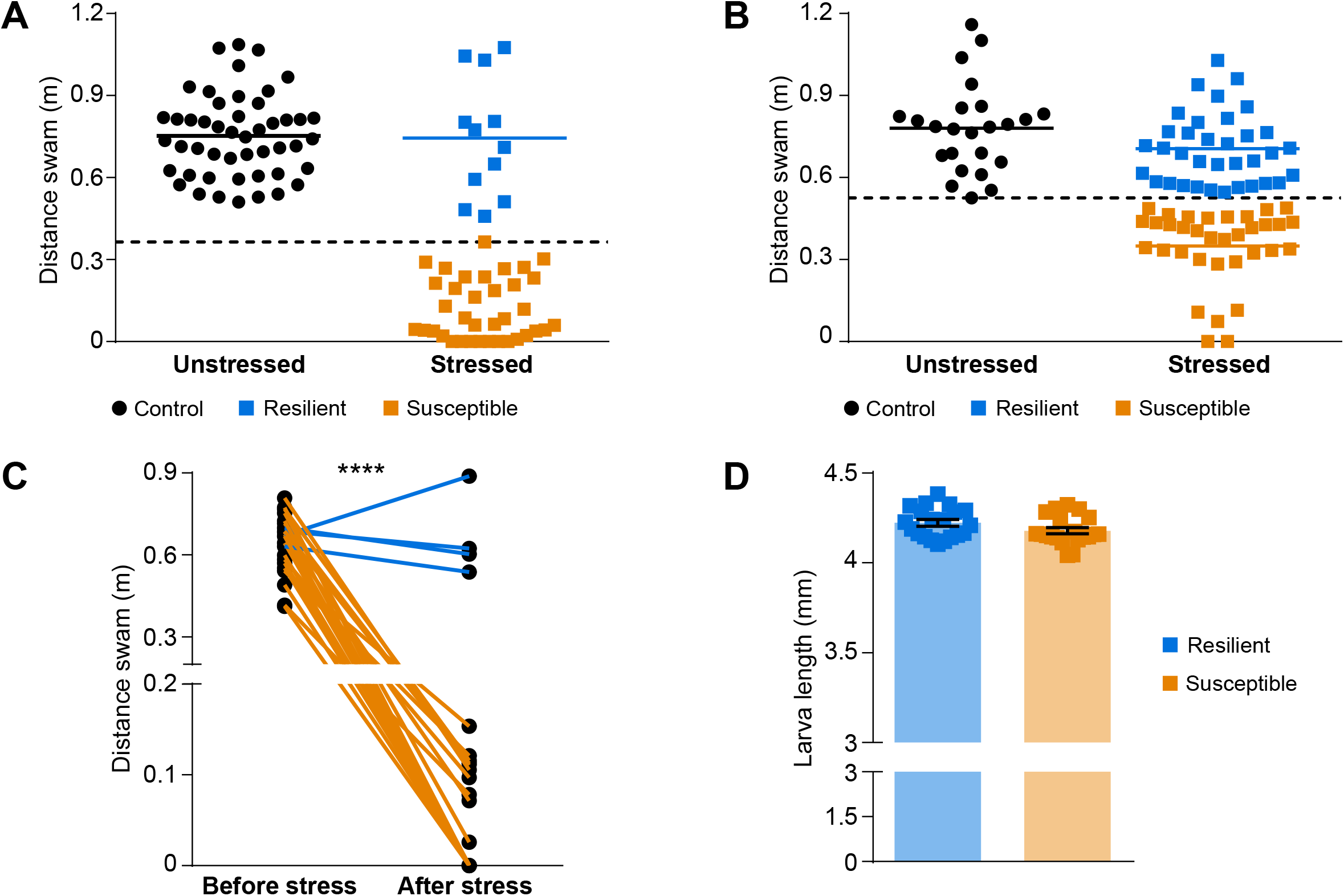
(**A**) 6 dpf larvae from a different wildtype strain, TL, were subjected to the two-step resilience assay. Larvae were subjected to osmotic stress for 20 minutes followed by recording their behaviour for one hour under mild stress (dark novel environment). The dot plot represents the first 5 minutes of the recovery from stress. (**B**) 6 dpf AB larvae were subjected to netting stress by removing out of water for 10 minutes. Behaviour was monitored under mild stress for one hour and the first 5 minutes are represented in the dot plot. (**C**) Basal locomotion of 6 dpf larvae was monitored in the dark (Before stress). Larvae were subjected to 20 minutes osmotic stress and behaviour was monitored following the stress (After stress). (**D**) Larvae were distinguished as resilient or susceptible from the two-step resilience assay and larval length was measured. Data presented as mean±SEM, **** p<0.0001

**Supplementary figure S2:**
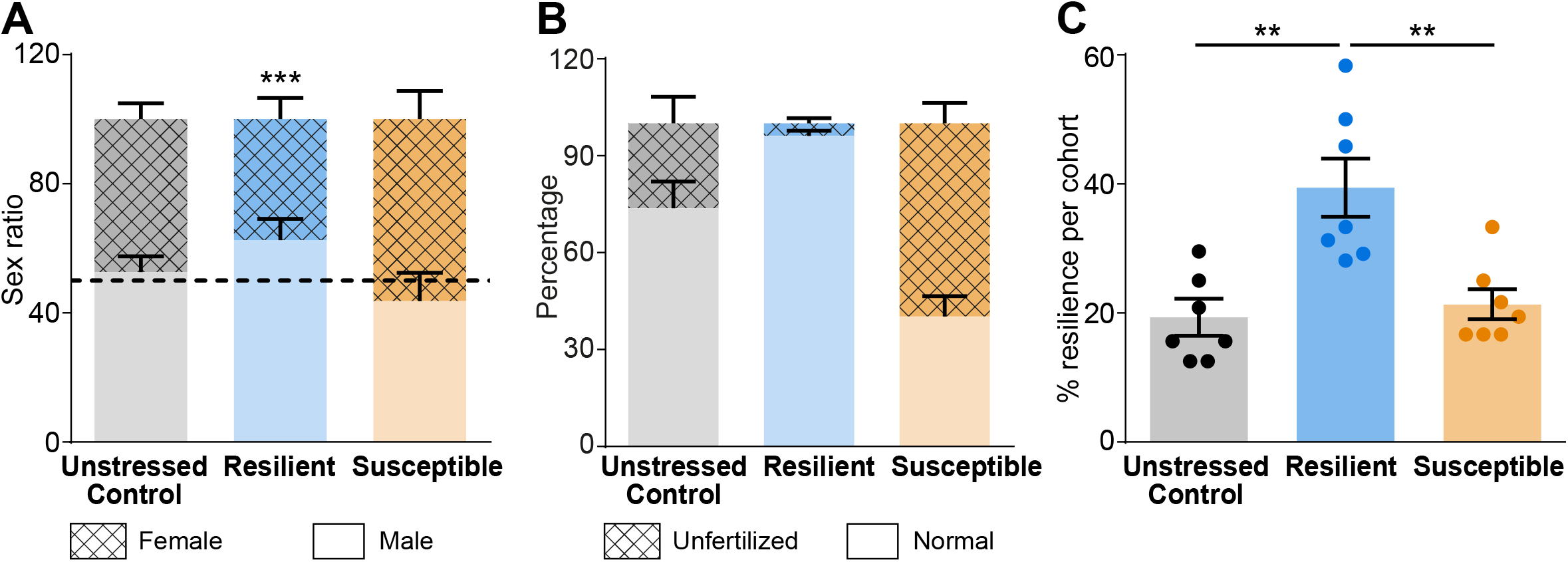
(**A**) Unstressed control, resilient and susceptible larvae were grown to adulthood and the proportion of males and females was quantified in each group. (**B**) Unstressed control, resilient and susceptible groups were incrossed. The percentage of eggs that were fertilized vs. those that died within the first 2 days (unfertilized) was quantified. (**C**) Percentage of resilience in the progeny of the unstressed control, resilient and susceptible fish was quantified (Figure 2E). Data compiled from multiple experiments is presented. Data presented as mean±SEM, ***p<0.001, **p<0.01

**Supplementary figure S3:**
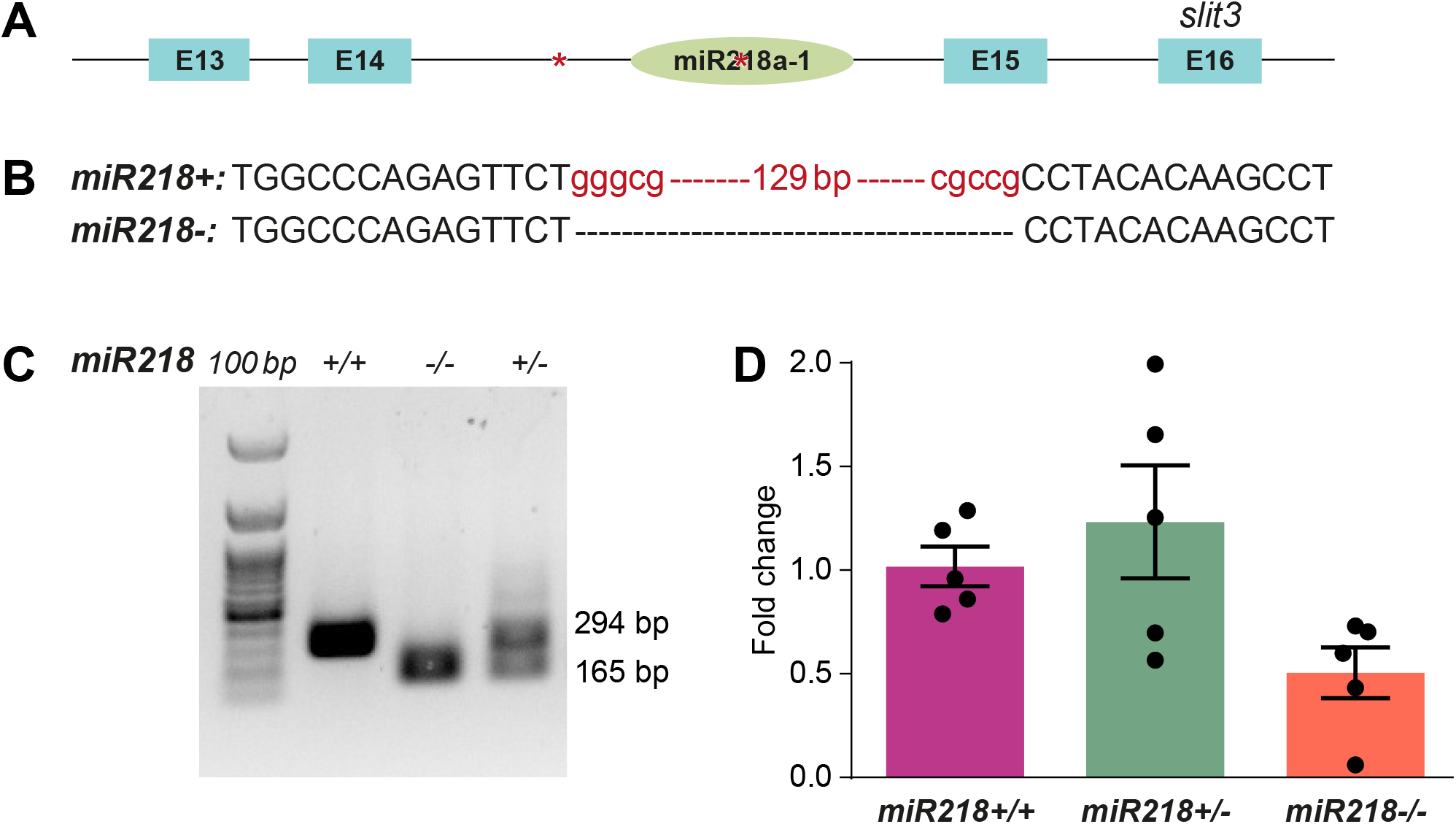
(**A**) Structure of the *slit3* gene showing the presence of *miR218a-1* in intron 14. Exons are marked as boxes, CRISPR sites depicted with stars. (**B**) Scheme showing the nature of deletion and flanking sequences in *miR218a-1*. (**C**) Genotyping of *miR218a-1* mutant from whole larval genomic DNA by PCR. (**D**) Expression analysis of *c5* and *c9* in wildtype, heterozygous and homozygous mutants by qRT-PCR. Data presented as mean±SEM, * p<0.05

**Supplementary figure S4:**
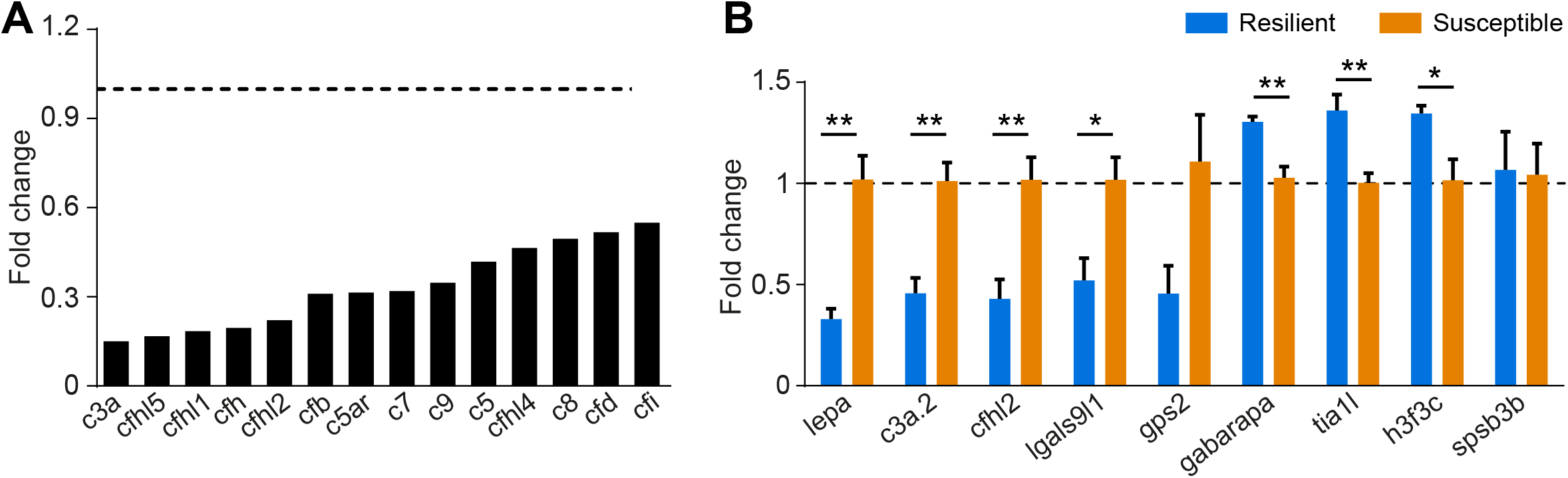
(**A**) Stress-responsive expression of various complement factors in resilient larvae from the RNA-sequencing analysis. (**B**) Gene expression was analyzed in resilient and susceptible larvae 30 minutes after 5 minute netting stress. (**C**) List of up- and downregulated genes was analyzed using a single cell RNA-sequencing atlas and the clusters where the genes are expressed was scored. The histogram shows the number of downregulated (blue) or upregulated (yellow) genes that are expressed in each cell type. Data presented as mean±SEM, **p<0.01, *p<0.05

**Supplementary figure S5:**
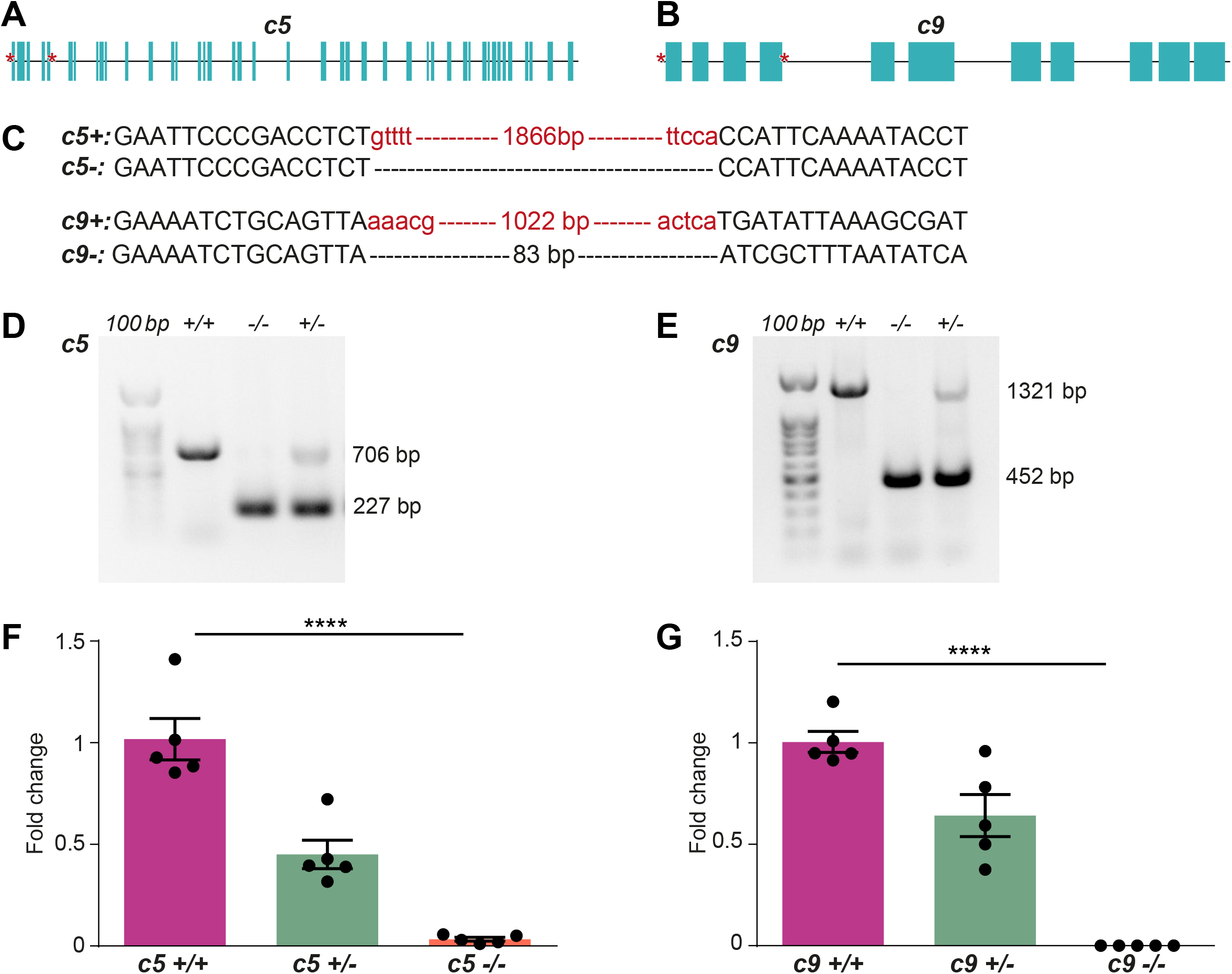
(**A, B**) Structure of the *c5* (A) and *c9* (B) genes. Exons are marked as boxes, CRISPR sites depicted with stars. (**C**) Scheme showing the nature of deletion and flanking sequences in *c5* and *c9* genes. (**D,E**) Genotyping of *c5* and *c9* mutants from whole larval genomic DNA by PCR. For genotyping *c5* mutants, a combination of primers flanking the deleted region and an internal primer were used. Mutants show amplification with the flanking primers, wildtype allele is amplified by the internal primer (D). For genotyping the c9 mutants, primers flanking the deleted region were used (E). (**F,G**) Expression analysis of *c5* and *c9* in wildtype, heterozygous and homozygous mutants by qRT-PCR. Data presented as mean±SEM, **** p<0.0001

**Supplementary figure S6:**
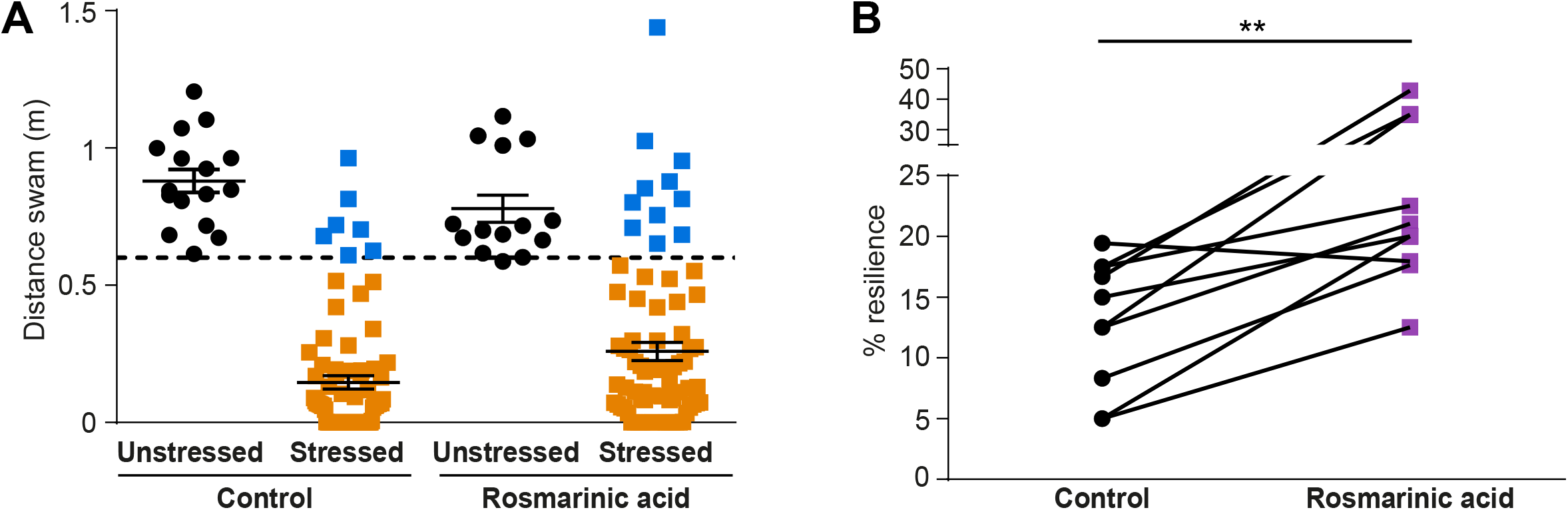
Larvae were treated with rosmarinic acid, a C3 and C5 convertase antagonist overnight and the effect on resilience was checked. In comparison to the control group, the treated group showed greater number of resilient fish (A-Compare blue squares between control and rosmarinic acid groups; B-data compiled from multiple experiments). Data presented as mean±SEM, ** p<0.01

**Supplementary table S1:** Processed RNA-sequencing data from resilient and susceptible larvae at basal and stressed conditions. Log fold change, adjusted p value, direction of change and whether statistically significant are presented for the six different comparisons between the four groups.

**Supplementary table S2:** Functional enrichment analysis of differentially expressed genes with the list of gene ontology terms performed using Metascape and Enrichr. The lists of downregulated and upregulated genes were separately analyzed using both tools.

**Supplementary table S3:** Cell type localization of differentially expressed genes. The cell types expressing each of the significantly down- and upregulated genes was analyzed using a scRNA-seq atlas. Members of the complement pathway are highlighted in green.

### List of oligonucleotides used in this study

#### Guide RNAs for CRISPR mutagenesis

C5: CRISPR 1: GCTTCATGGGAATAAAACAGAGG

CRISPR 2: ATACATACCTTATAGTGACATGG

C9: CRISPR 1: GAAAATCTGCAGTTAAAACGAGG

CRISPR 2: GATGTCATAAGGTCACCCTGCGG

miR218: CRISPR 1: TGTGAGGCTTGTGTAGGCGGCGG

CRISPR 2: TGCTGGCCCAGAGTTCTGGGCGG

#### Primers for genotyping

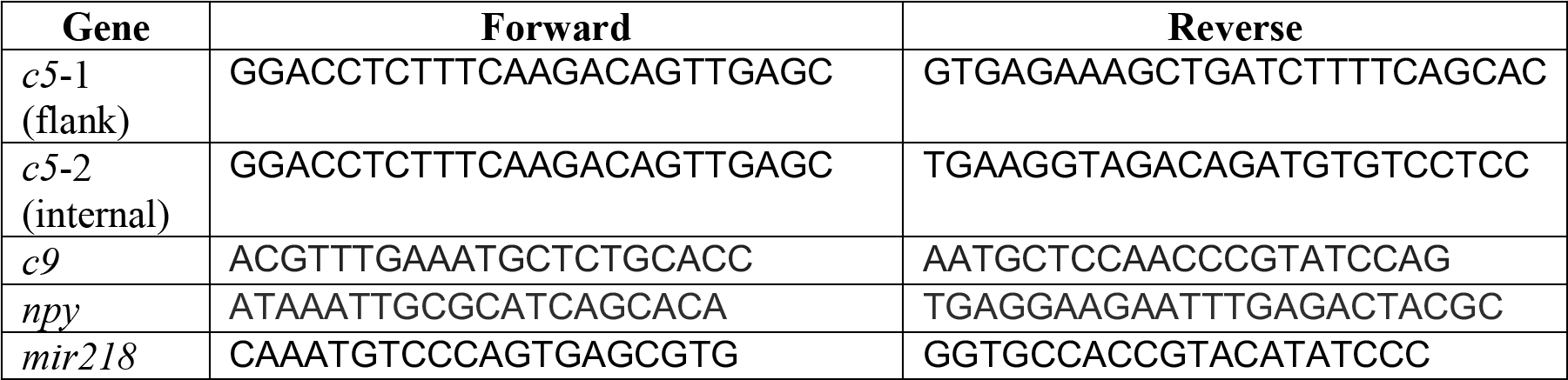

#### Primers for qRT-PCR

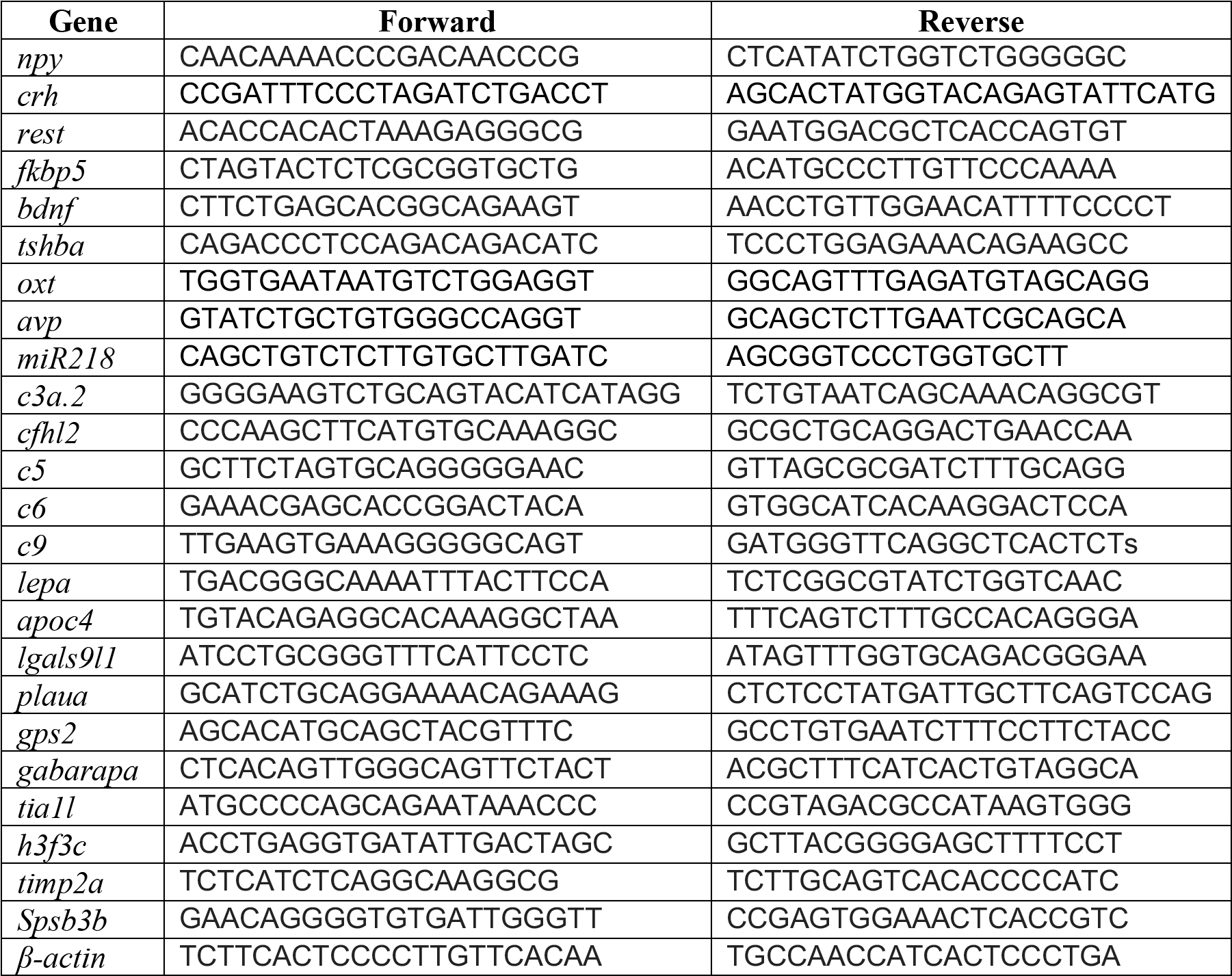

